# Comparative Transcriptomic Analysis of Gene Expression Patterns in Neutrophils from Rheumatoid Arthritis and COVID-19

**DOI:** 10.64898/2026.11.03.764103

**Authors:** Sakshi Nawani, Mansaf Alam

## Abstract

**Background:** Rheumatoid arthritis (RA) is an autoimmune disorder characterised by chronic inflammation and immune dysregulation, and neutrophils play a key role in its progression. COVID-19 also involves neutrophil hyperactivation that contributes to severe immune responses. Emerging evidence links COVID-19 to increased incidence or exacerbation of RA, potentially through overlapping immune mechanisms.

**Objective:** To compare neutrophil transcriptomic profiles in RA and COVID-19 and identify shared immune pathways and disease-specific features.

**Methods:** Publicly available neutrophil RNA-seq count data from the Gene Expression Omnibus were analysed: RA versus healthy controls (GSE116899) and COVID-19 versus healthy controls (GSE212041). A second COVID-19 comparison (severe versus convalescent) was assembled from GSE272381 and GSE227116. Differential expression was analysed with DESeq2, pathway enrichment with fgsea (Hallmark gene sets) and topGO (Gene Ontology biological process and molecular function). Differentially expressed genes (DEGs) were compared between diseases by direction of change.

**Results:** In RA neutrophils, 11,355 DEGs were identified, of which 90 had an absolute log2 fold change of at least 2 with p<0.05 (70 upregulated, 20 downregulated). Twenty-four Hallmark gene sets were positively enriched at adjusted p<0.05, led by interferon alpha and interferon gamma responses, and Gene Ontology analysis highlighted type I interferon, antiviral and double-stranded RNA terms. In COVID-19 neutrophils, 420 DEGs were identified (37 upregulated, 383 downregulated), and no Hallmark gene set reached adjusted p<0.05. Of the 90 RA DEGs, 46 changed in the same direction in COVID-19 without reaching significance, and 27 changed significantly in the opposite direction, including CAMK1G, SIGLEC1, OAS1 and OAS3. In the second COVID-19 comparison, several of these genes, including OAS1, OAS3 and SIGLEC1, changed in the same direction as in RA.

**Conclusions:** RA neutrophils show a transcriptional programme enriched for interferon and antiviral-response pathways. Part of the RA signature is shared with COVID-19 neutrophils, while a subset of genes diverges, and this divergence depended on the COVID-19 comparison used. These findings are hypothesis-generating and provide candidate genes for experimental validation.

## 1. Introduction

Autoimmune diseases such as systemic lupus erythematosus and rheumatoid arthritis (RA) arise when the immune system mistakenly attacks self-antigens, often causing chronic inflammation and tissue damage. Viral infections are recognised environmental triggers for the onset or exacerbation of autoimmunity, through mechanisms including molecular mimicry, bystander activation and epitope spreading; hepatitis B virus, parvovirus B19 and herpes viruses have historically been linked to these processes [1], [2]. Type I interferons (IFN-I) have a dual role: excessive activity contributes to autoimmune disease and correlates with severity in autoimmune patients, whereas IFN-I also controls viral replication and supports antigen presentation. Viral resistance and autoimmune susceptibility have been reported to share transcriptomic and genomic signatures [3].

Neutrophils, the most abundant white blood cells and a first line of defence, are now recognised as functionally complex cells that fight pathogens and also regulate adaptive immune responses. Their cytokines, chemokines and growth factors balance host protection against tissue damage [4].

RA affects approximately 1–2% of the global population and is the most common form of chronic inflammatory arthritis, characterised by synovial inflammation leading to joint destruction. Autoantibodies, including rheumatoid factors and anti-citrullinated protein antibodies (ACPAs), and elevated C-reactive protein, IL-6 and TNF-α are central features [5]. Neutrophils are abundant in synovial fluid and release elastase, cathepsin G and myeloperoxidase; dysregulated neutrophil extracellular trap (NET) formation releases citrullinated proteins that can act as autoantigens driving ACPA production [5], [6].

COVID-19 is caused by SARS-CoV-2, which enters cells via ACE2. Severe disease involves a dysregulated inflammatory response (“cytokine storm”) that can lead to acute respiratory distress syndrome and multi-organ failure, with neutrophilia, an increased neutrophil-to-lymphocyte ratio, and neutrophil release of reactive oxygen species and proteases. SARS-CoV-2 can also antagonise interferon signalling, delaying antiviral responses [7].

A large cohort study reported that COVID-19 is associated with an increased risk of developing RA and other autoimmune disorders [8], [9]. RA and COVID-19 share features including exaggerated immune responses with a central role for neutrophils, and involvement of the type I interferon response [7], [8], [10], [11]. Transcriptomic studies of RA have historically focused on peripheral blood mononuclear cells, with a shift towards specific subsets such as CD4+ T cells and neutrophils [12], while COVID-19 transcriptomic studies have reported altered cytokine-signalling and antiviral pathways [13], [14]. However, studies comparing neutrophil-specific transcriptomes of RA and COVID-19 are scarce.

Here, we compared neutrophil RNA-seq profiles from RA and COVID-19 using public datasets. Our objectives were to (i) compare neutrophil transcriptomic profiles in the two diseases, (ii) identify DEGs and their roles in shared pathways, such as type I interferon responses, and in disease-specific features, (iii) examine genes including CAMK1G, SIGLEC1, OAS1 and OAS3 that differed between the diseases, and (iv) provide a foundation for further study of the link between COVID-19 and RA.

## 2. Materials and Methods

### 2.1 Datasets

Neutrophil RNA-seq feature (read) count data were retrieved from the Gene Expression Omnibus (GEO) [15] (Table 1).

**Table 1.** Datasets analysed.

| Comparison | GEO accession | Public on | Description |
|---|---|---|---|
| RA vs healthy | GSE116899 | 11 Jul 2018 | Synovial fluid and peripheral blood neutrophils from RA patients and matched healthy controls (GEO record: 7 RA patients and 5 healthy controls). The analysed count matrix comprised 30 samples (24 RA, 6 healthy), including technical replicates (the same library sequenced on two lanes). |
| COVID-19 vs healthy (cohort 1) | GSE212041 | 28 Aug 2022 | Circulating neutrophils from hospitalised COVID-19 patients sampled on the first day of admission (WHO outcome scale category A5; discharged without admission within 28 days) and 8 healthy controls: 70 samples (62 COVID-19, 8 healthy) [16], [17]. |
| Severe COVID-19 vs healthy | GSE272381 | 23 Jul 2024 | Neutrophils and low-density granulocytes from COVID-19 patients. |
| Convalescent vs healthy | GSE227116 | 29 Mar 2023 | Blood samples from convalescent COVID-19 patients and healthy donors. |

To reduce errors arising from differences in methodology, patient sampling or infection stage, data from independent studies were used as a second COVID-19 comparison. The severe-versus-healthy and convalescent-versus-healthy neutrophil datasets were merged to obtain a severe-versus-convalescent comparison (15 severe and 66 convalescent samples), referred to below as cohort 2 (labelled “acute vs convalescent” in the figures).

### 2.2 Differential expression analysis

Raw count matrices (genes as rows, samples as columns) were analysed in R using DESeq2, which models counts with a negative binomial distribution, performs internal library-size normalisation, and estimates log2 fold changes, p-values and adjusted p-values [18], [19]. Analyses were run for three comparisons: RA vs healthy, COVID-19 vs healthy, and severe vs convalescent. DEGs were defined as genes with an absolute log2 fold change of at least 2 and p<0.05. Volcano plots were generated for each comparison.

### 2.3 Functional and pathway analysis

Gene set enrichment analysis (GSEA) was run with the fgsea R package, using Hallmark gene sets obtained through msigdbr [20], [21]. Gene Ontology (GO) enrichment of DEGs for biological process (BP) and molecular function (MF) terms was run with topGO using org.Hs.eg.db annotation [22]. Results were displayed as enrichment plots, bar plots and dot plots.

### 2.4 Comparative analysis

DEGs from RA vs healthy and COVID-19 vs healthy were each derived against their own healthy controls and then compared, which avoids batch effects between laboratories and the biases of batch-correction tools. Overlapping DEGs were classified as changing in the same or in the opposite direction. The cohort 2 DEGs were then compared with the RA and cohort 1 results, and GO enrichment was applied to the discordant gene sets.

## 3. Results

### 3.1 Differential expression and pathways in RA neutrophils

Differential expression analysis of RA versus healthy neutrophils identified 11,355 DEGs, of which 90 had an absolute log2 fold change of at least 2 with p<0.05 (70 upregulated, 20 downregulated) (Fig. 1). Genes in the output snippet (Supplementary Fig. S1) include VCAN, TRAF1, NEBL, SIGLEC1, OAS1, RPH3A, KCNH4, CREM, TLDC2, SFRP1, SEPT4, OAS3, CCL20, FN1, CCRL2, ADORA3 and P4HA1, all with positive log2 fold changes.

**Figure 1.**
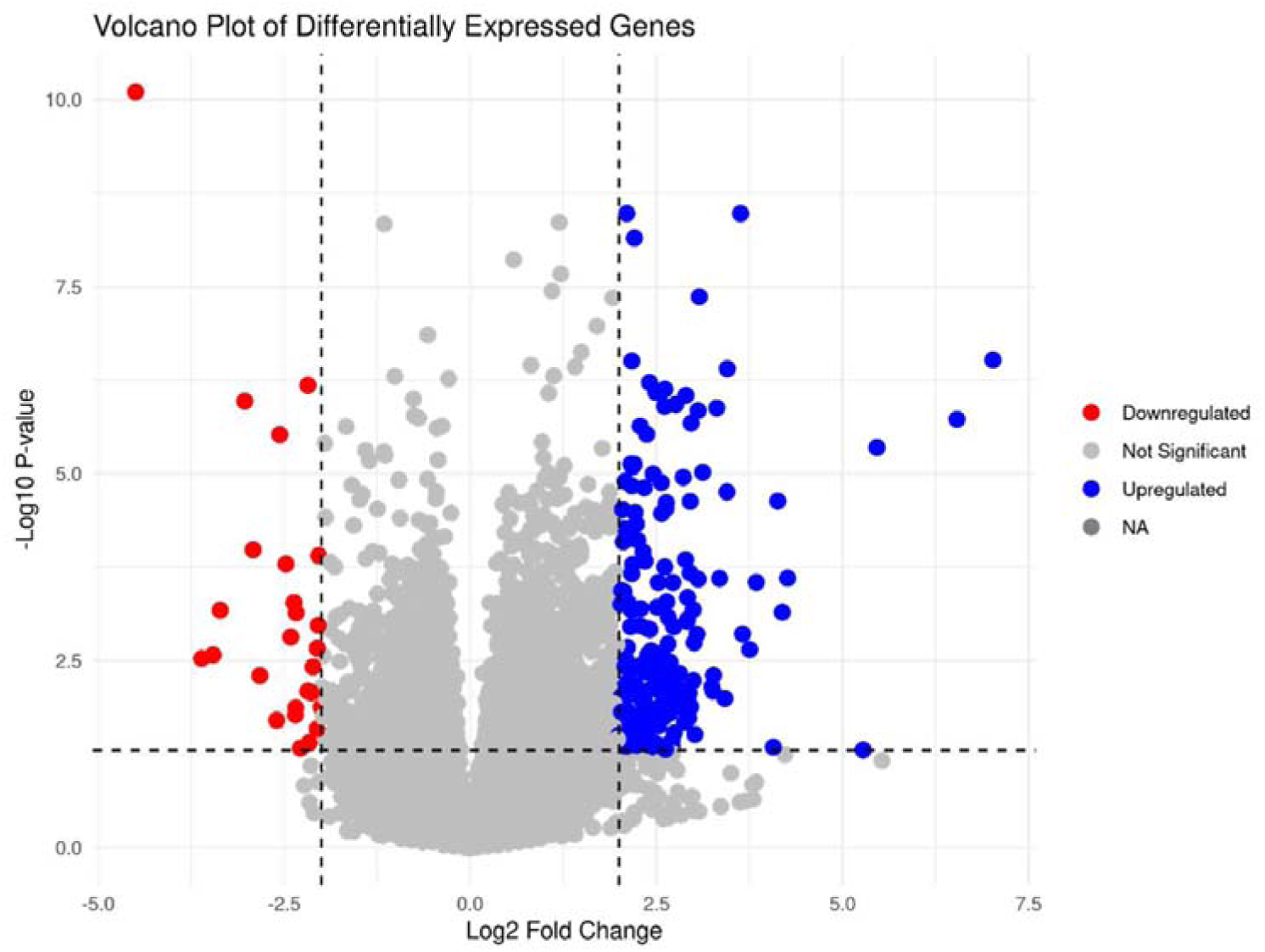
Differentially expressed genes in RA versus healthy control neutrophils. Volcano plot of all genes (GSE116899); red, downregulated; blue, upregulated; grey, not significant. Dashed lines mark the significance and fold-change thresholds.

GSEA displayed 34 Hallmark gene sets at an FDR-adjusted p<0.25 (Fig. 2A). Of these, 24 positively enriched sets and 2 negatively enriched sets (E2F targets and G2M checkpoint) had adjusted p<0.05. Interferon alpha response and interferon gamma response had the highest normalised enrichment scores, followed by epithelial-mesenchymal transition, inflammatory response and TNF-α signalling via NF-κB. Enrichment plots for the two top pathways by p-value, interferon alpha response and inflammatory response, are shown in Fig. 2B.

**Figure 2.**
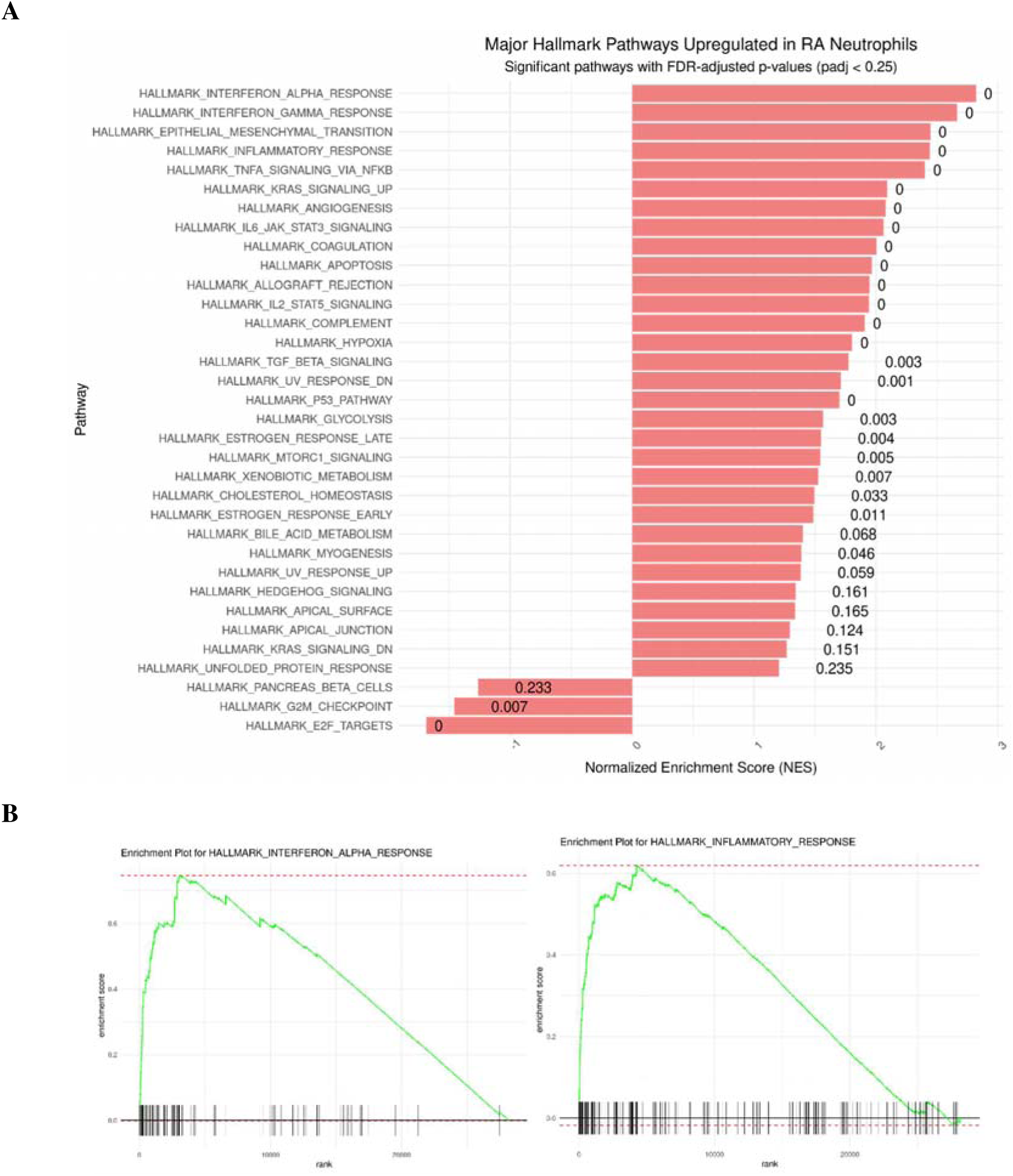
Hallmark gene set enrichment in RA neutrophils compared with healthy control neutrophils. (A) Normalised enrichment scores (NES) of Hallmark gene sets displayed at FDR-adjusted p<0.25; adjusted p-values are labelled. (B) Enrichment plots for the Hallmark interferon alpha response and inflammatory response gene sets, the two top pathways by p-value.

GO enrichment of the 90 DEGs showed enriched BP and MF terms (Fig. 3). Top BP terms included negative regulation of viral genome replication, negative regulation of IP-10 production, the IL-27-mediated signalling pathway (p = 9.80e-07), regulation of ribonuclease activity (p = 2.30e-06), antiviral innate immune response (p = 1.30e-05) and negative regulation of type I interferon-mediated signalling pathway (p = 3.10e-05); defense response to virus (p = 1.23e-03) and cellular response to type I interferon (p = 5.78e-03) were also among the top 25. The most enriched MF term was 2′-5′-oligoadenylate synthetase activity, followed by double-stranded RNA binding. Together, these results indicate that RA neutrophils show a broad host-defence response reminiscent of antiviral immunity, even though RA is not generally considered a pathogen-driven disease.

**Figure 3.**
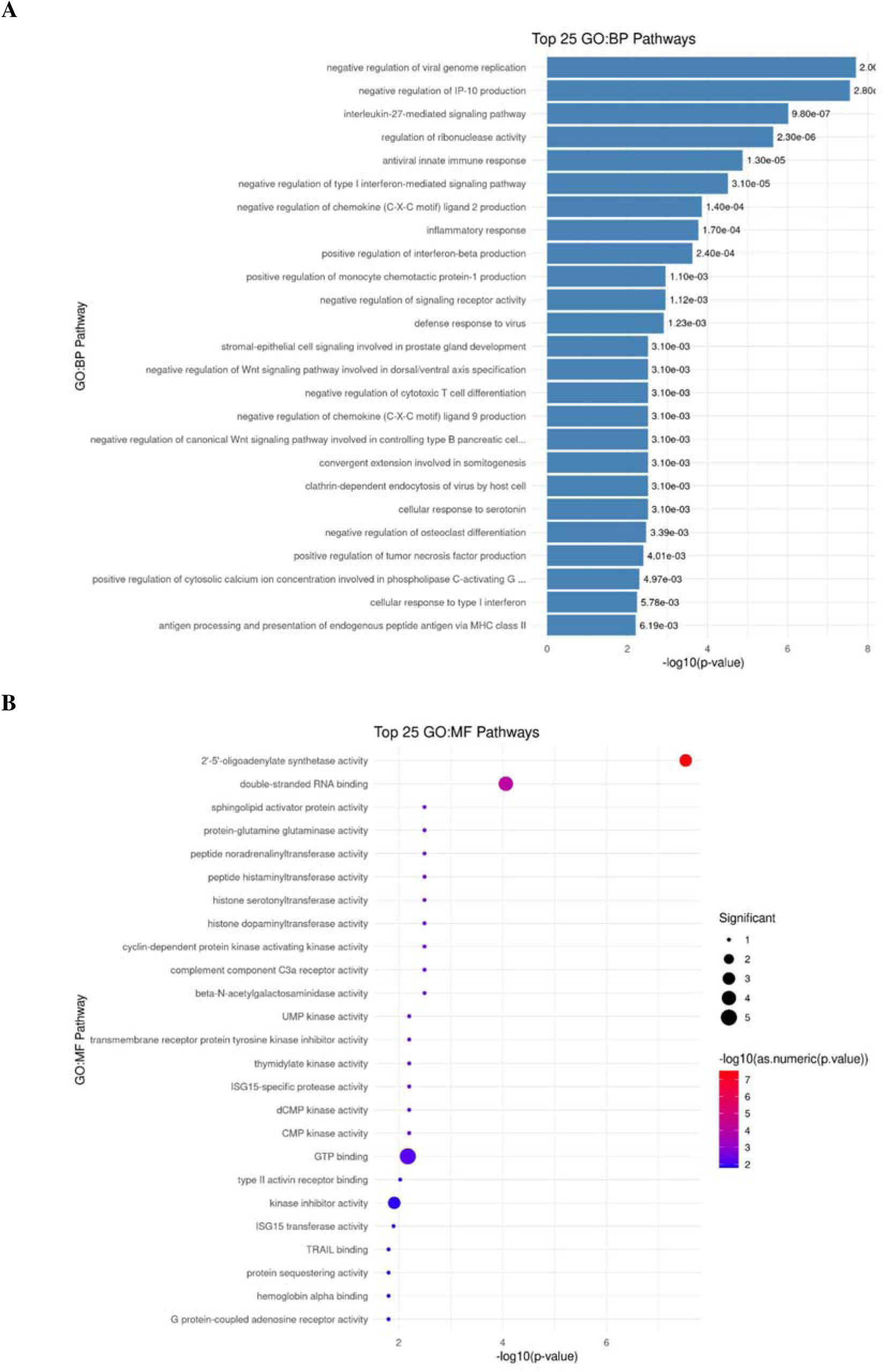
Gene Ontology enrichment of the 90 DEGs in RA neutrophils. (A) Top 25 biological process (BP) terms with p-values. (B) Top 25 molecular function (MF) terms; point colour indicates −log10(p-value) and point size the number of significant genes.

### 3.2 Differential expression and pathways in COVID-19 neutrophils (cohort 1)

In COVID-19 versus healthy neutrophils, 420 DEGs were identified at an absolute log2 fold change of at least 2 and p<0.05 (37 upregulated, 383 downregulated) (Fig. 4). Genes in the output snippet (Supplementary Fig. S2) include ETV7, LTF, TNFRSF17, DHRS9, CA12, DUSP13, CEACAM6, ADAMTS2, SIGLEC1, RPH3A, BIRC5, DSP and BPI, all with negative log2 fold changes.

**Figure 4.**
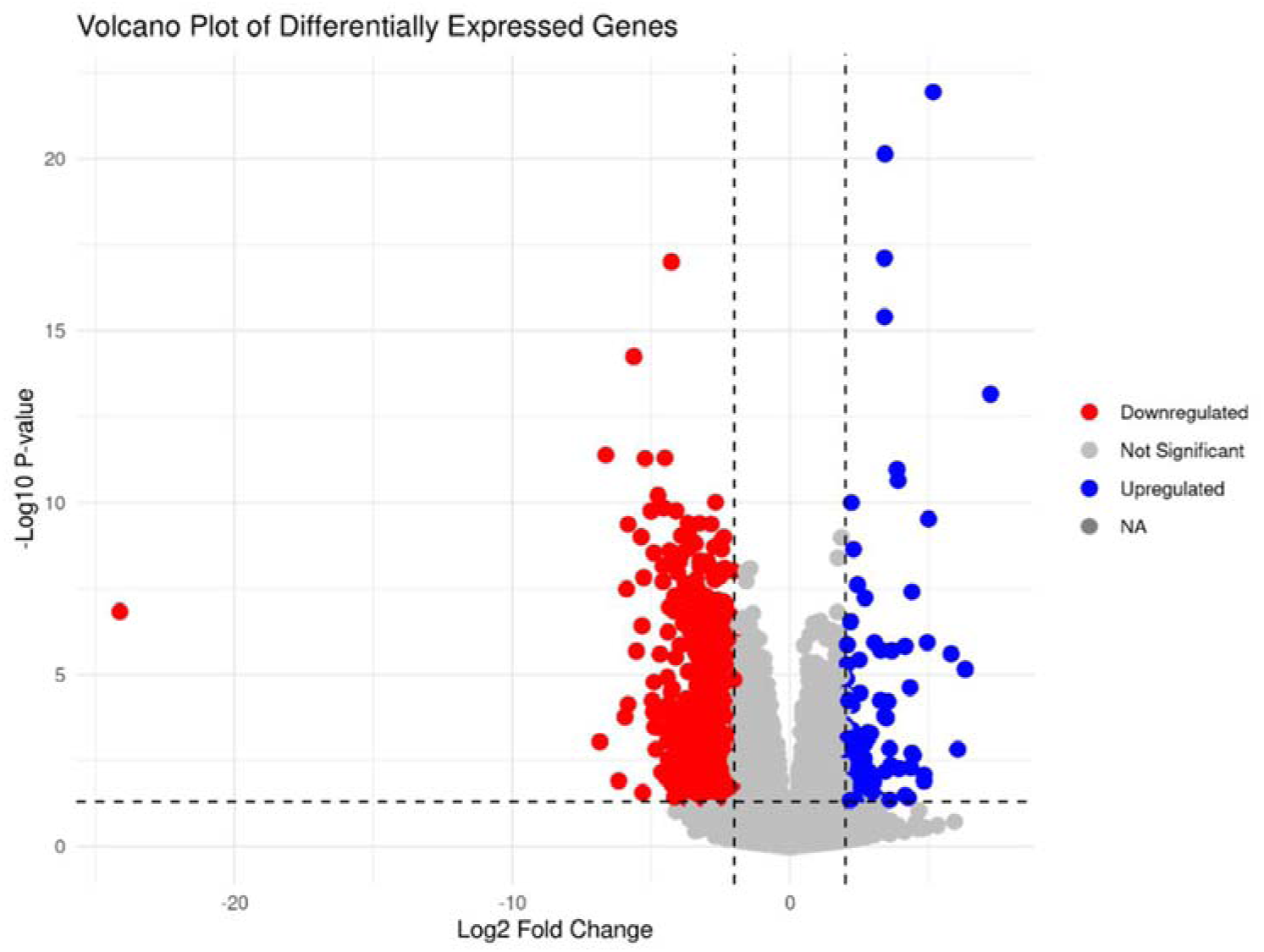
Differentially expressed genes in COVID-19 versus healthy control neutrophils. Volcano plot of all genes (GSE212041); colours as in Fig. 1.

GSEA displayed 24 Hallmark gene sets at an adjusted p<0.25 (Fig. 5A): 15 with positive and 9 with negative enrichment scores. The lowest adjusted p-value displayed was 0.067, so none reached adjusted p<0.05. Twenty-one of the 24 gene sets were also among those displayed for RA (Fig. 2A), but the direction differed for several: interferon alpha response, interferon gamma response, glycolysis, hypoxia, IL-2/STAT5 signalling, mTORC1 signalling, apical junction and unfolded protein response had positive enrichment scores in RA and negative scores in COVID-19, whereas E2F targets showed the opposite pattern. The two top pathways shared with RA by p-value were KRAS signalling up and apoptosis (Fig. 5B).

**Figure 5.**
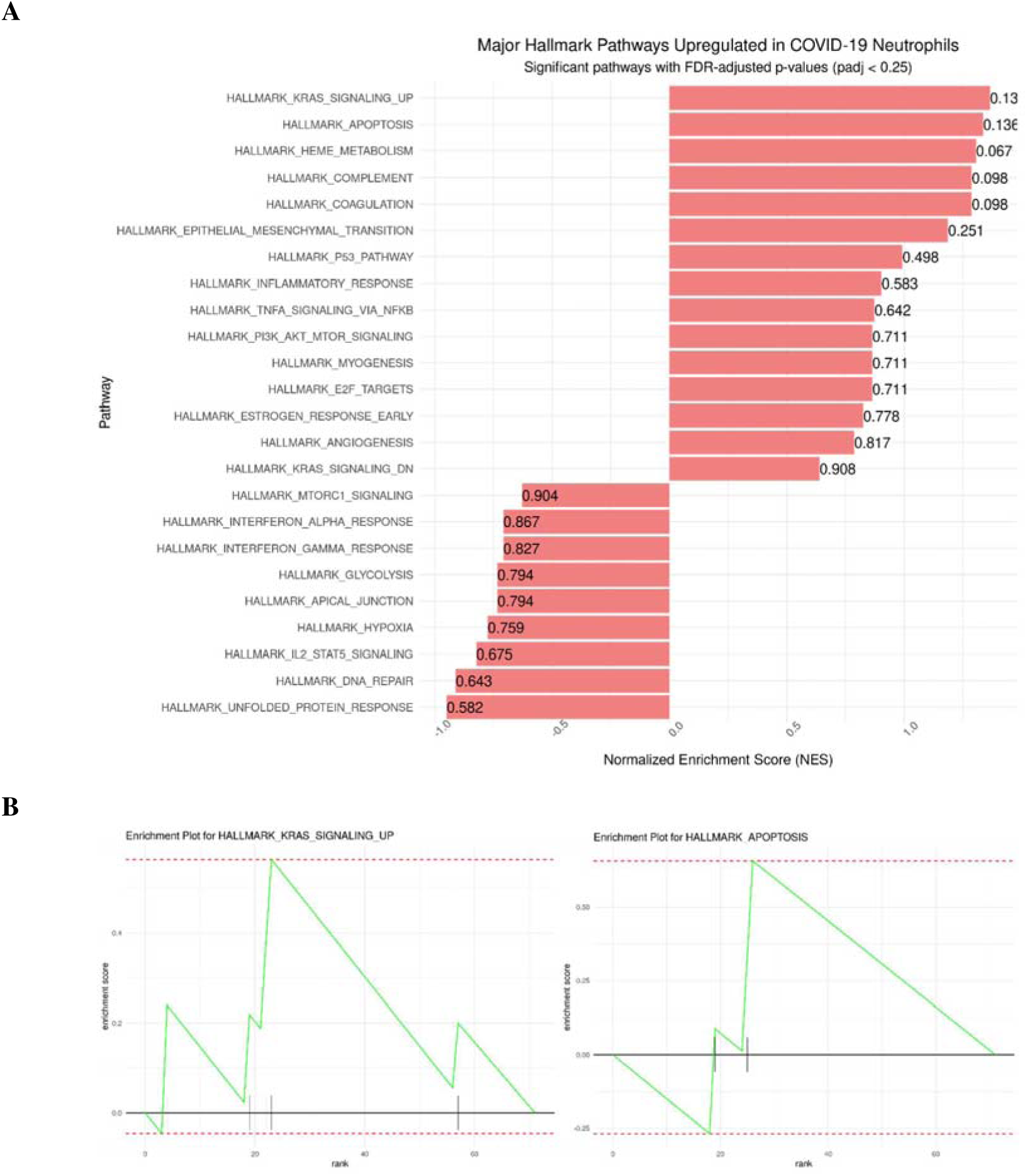
Hallmark gene set enrichment in COVID-19 neutrophils compared with healthy control neutrophils. (A) NES of Hallmark gene sets displayed at FDR-adjusted p<0.25; adjusted p-values are labelled. (B) Enrichment plots for the KRAS signalling up and apoptosis gene sets, the two top pathways by p-value shared with RA.

Top GO terms among the COVID-19 DEGs overlapped substantially with those in RA (Fig. 6): negative regulation of viral genome replication, defense response to virus (p = 2.60e-06), negative regulation of type I interferon-mediated signalling pathway (p = 1.40e-04), positive regulation of interferon-beta production, cellular response to type I interferon, regulation of ribonuclease activity, negative regulation of IP-10 production, and 2′-5′-oligoadenylate synthetase activity. Response to interferon-beta (p = 1.30e-04) and the MDA-5 signalling pathway (p = 1.09e-03) appeared among the top 25 BP terms in COVID-19 but not in the top 25 shown for RA.

**Figure 6.**
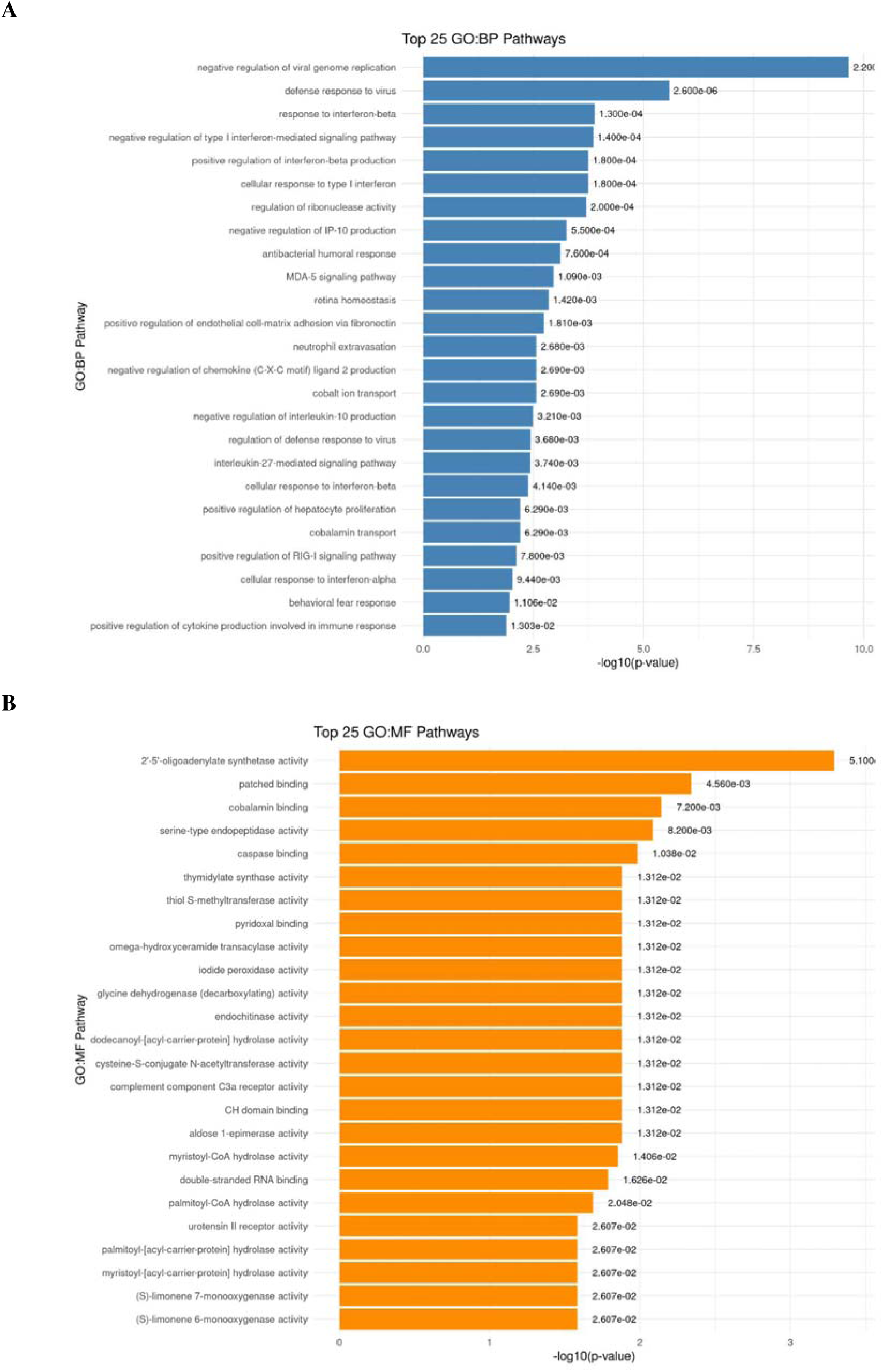
Gene Ontology enrichment of DEGs in COVID-19 neutrophils (cohort 1). (A) Top 25 BP terms with p-values. (B) Top 25 MF terms with p-values.

### 3.3 Direct comparison of RA and COVID-19 neutrophils

Of the 90 genes differentially expressed in RA, 46 changed in the same direction in COVID-19 but were not statistically significant there (p≥0.1, or p≥0.2 if fold change >2) (Fig. 7). In contrast, 27 genes were differentially expressed in opposite directions in the two diseases with statistical significance, being upregulated in RA and downregulated in COVID-19 (Fig. 8): XAF1, USP18, TCN2, SPATS2L, SIGLEC1, SEPT4, RTP4, RSAD2, RPH3A, RP11-561B11.1, RP11-466G12.3, RP11-408H1.3, RP11-288L9.1, RP11-24F11.2, RP11-177F15.1, OTOF, OAS3, OAS1, MAOA, IFIT1B, IFIT1, IFI44L, HESX1, DSP, CAMK1G, C3AR1 and C1QC.

**Figure 7.**
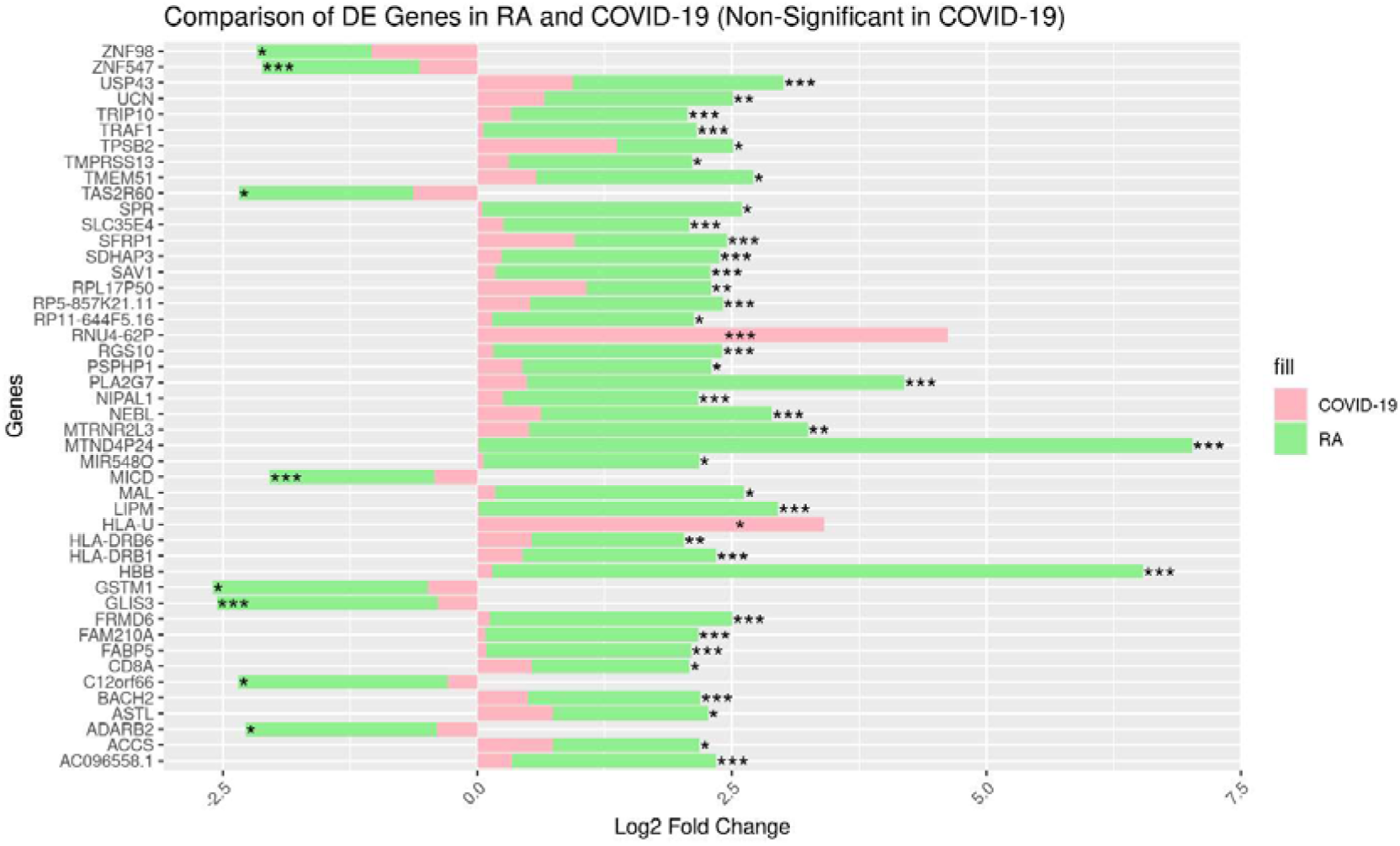
Genes differentially expressed in RA and changing in the same direction, but not significantly, in COVID-19. Log2 fold changes in RA (green) and COVID-19 (pink) for the 46 genes that are DE in RA (p<0.05) and expressed in the same direction, but not statistically significant (p≥0.1, or p≥0.2 if fold change >2), in COVID-19.

**Figure 8.**
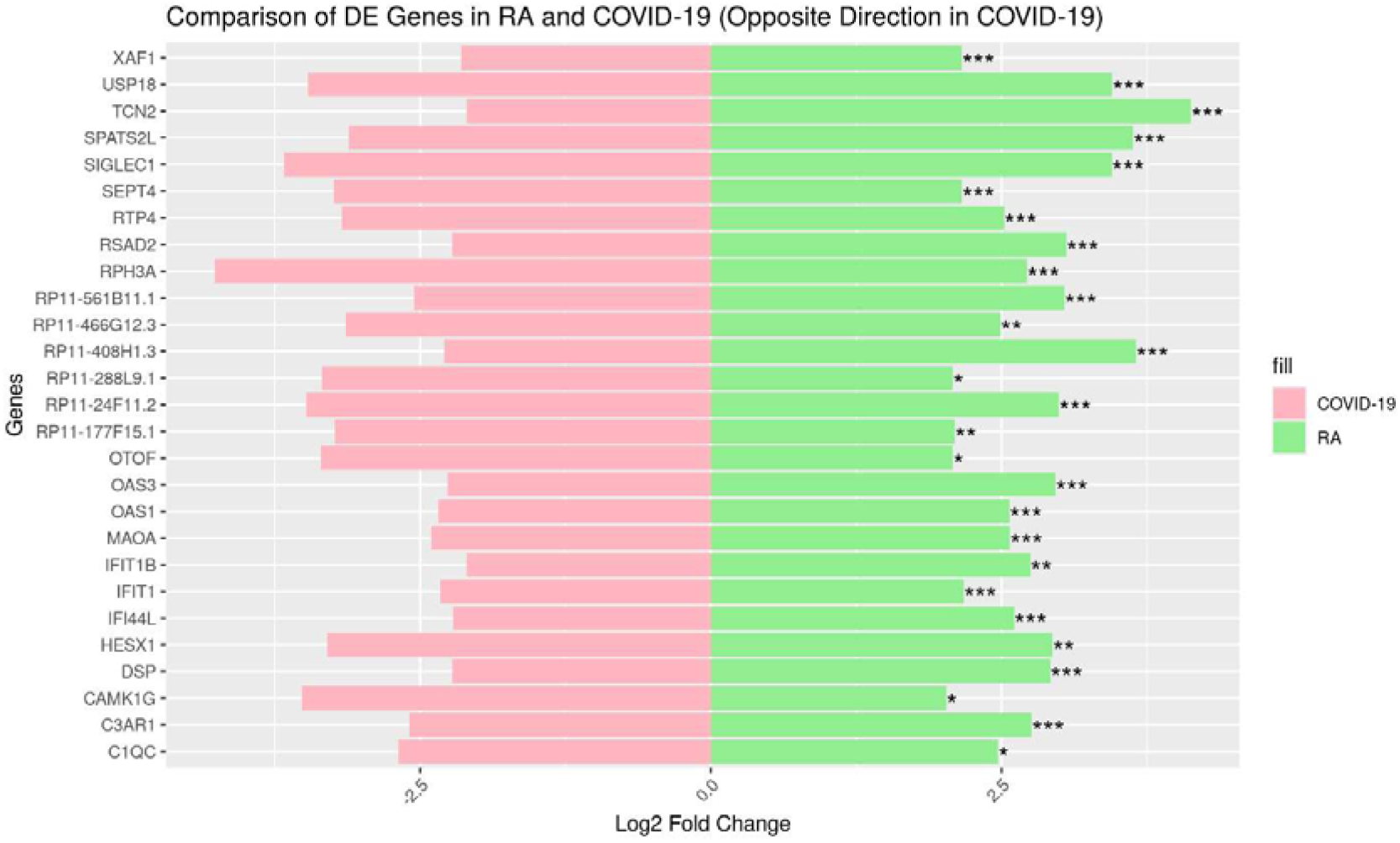
Genes differentially expressed in RA and changing in the opposite direction in COVID-19. Log2 fold changes in RA (green) and COVID-19 (pink) for the 27 DE genes in RA neutrophils (p<0.05) expressed in the opposite direction in COVID-19 neutrophils. *p<0.05, p<0.01,* **p<0.005.

GO analysis of the genes that differed between RA and COVID-19 showed enriched BP and MF terms (Fig. 9). Top BP terms were negative regulation of viral genome replication, cellular response to type I interferon (p = 6.90e-07), negative regulation of type I interferon-mediated signalling pathway (p = 1.00e-06), negative regulation of IP-10 production (p = 2.90e-06), negative regulation of chemokine (C-X-C motif) ligand 2 production (p = 1.50e-05), regulation of ribonuclease activity (p = 3.50e-05) and antiviral innate immune response (p = 1.00e-04). Top MF terms were 2′-5′-oligoadenylate synthetase activity, complement component C3a receptor activity (p = 1.10e-03), sepiapterin reductase activity (p = 1.90e-03), glutamic-type peptidase activity (p = 1.90e-03), monoamine oxidase activity (p = 3.30e-03) and ISG15-specific protease activity (p = 3.70e-03).

**Figure 9.**
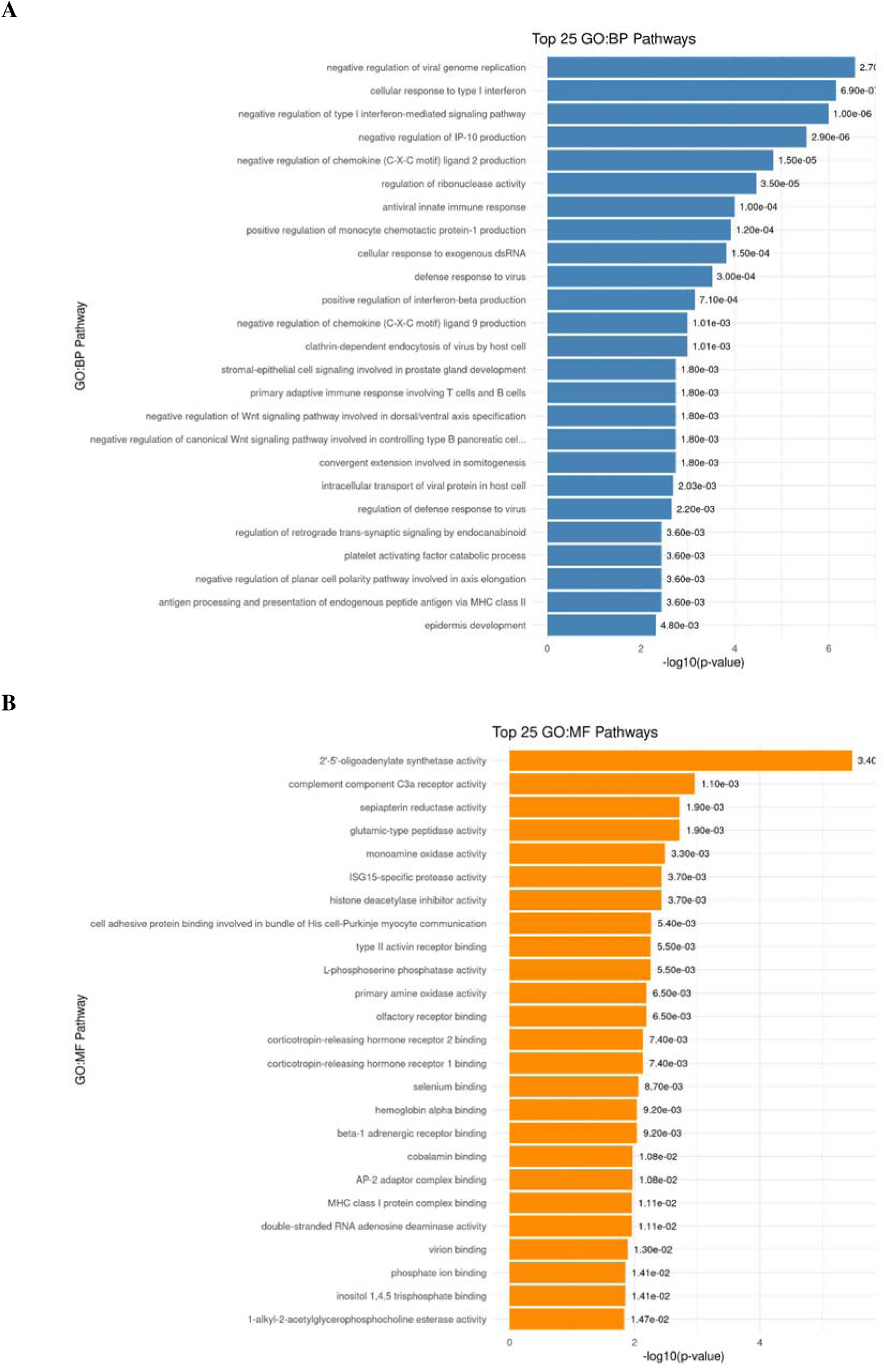
Gene Ontology enrichment of genes differing between RA and COVID-19 neutrophils. (A) Top 25 BP terms and (B) top 25 MF terms for the combined same- and opposite-direction gene sets, with p-values.

Within the pathway negative regulation of type I interferon-mediated signalling, USP18, TRIM25, TNFAIP3, TANK, SOCS3, SOCS1, ISG15, IRF2 and IFI44 were all upregulated in RA; in COVID-19, all were downregulated except SOCS3, which showed a small increase (Fig. 10).

**Figure 10.**
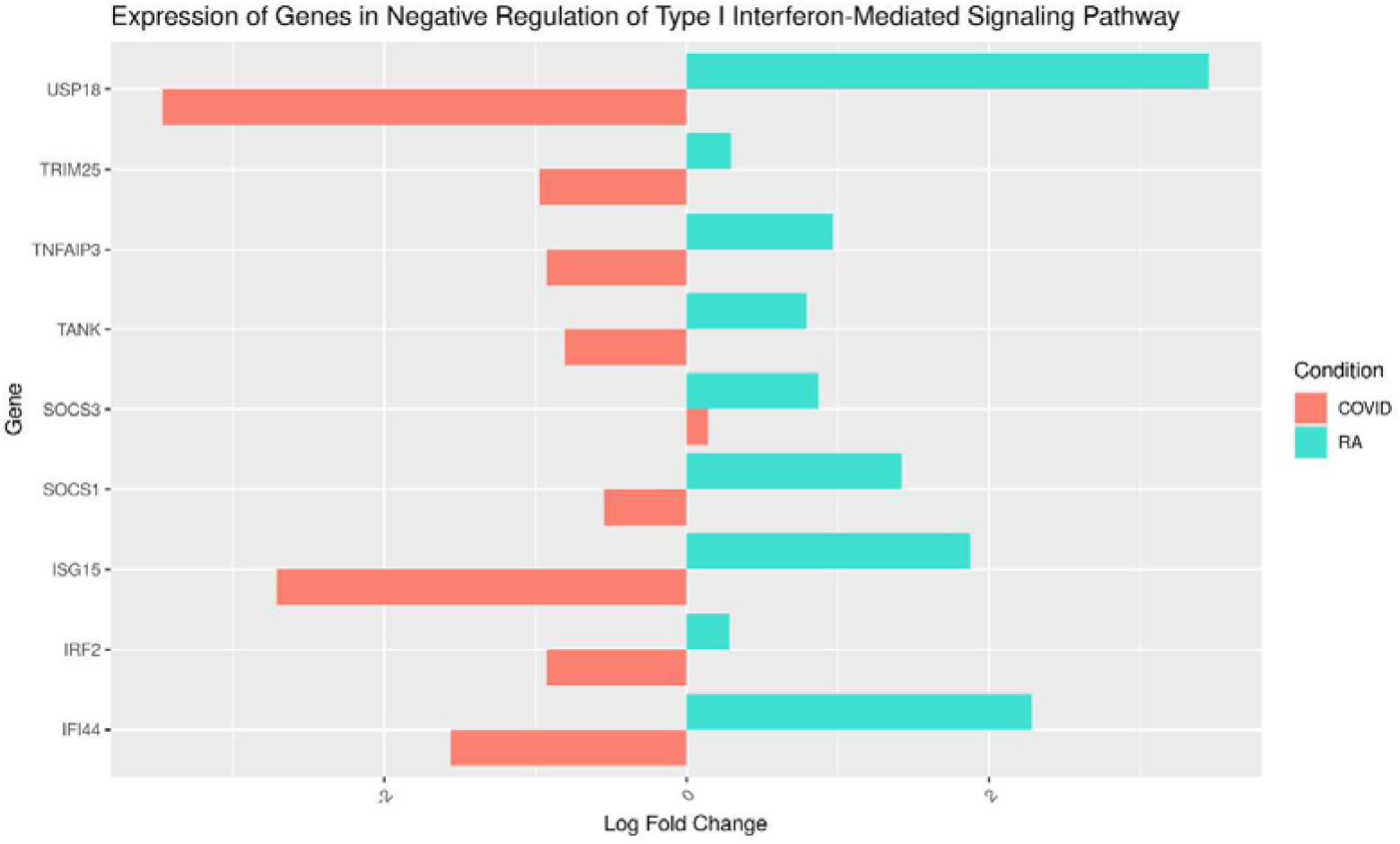
Expression of genes in the negative regulation of type I interferon-mediated signalling pathway. Log fold change in RA (turquoise) and COVID-19 (red).

### 3.4 Comparison with a second COVID-19 cohort

The RA DEGs were also examined in cohort 2 (acute versus convalescent COVID-19). Fig. 11A shows 36 genes with changes opposite to RA and Fig. 11B shows 25 genes changing in the same direction as RA. Thirteen genes changed in the same direction in RA, cohort 1 and cohort 2 (Fig. 11C): TYMS, RECQL4, PI3, NFXL1, MT-ND6, MT-CO3, LIMD1-AS1, KIR2DS4, IGLV2-14, IGHV3-15, HIST1H2AG, HCG4P5 and ANKRD18A. Thirty genes were upregulated in RA and downregulated in both COVID-19 comparisons (Fig. 11D), for example YPEL4, TCN2, PRTN3, PERP, IL4I1, IFIT1B, ICOS, EMP1, DUSP4, DSP, CCR6 and ANG.

**Figure 11.**
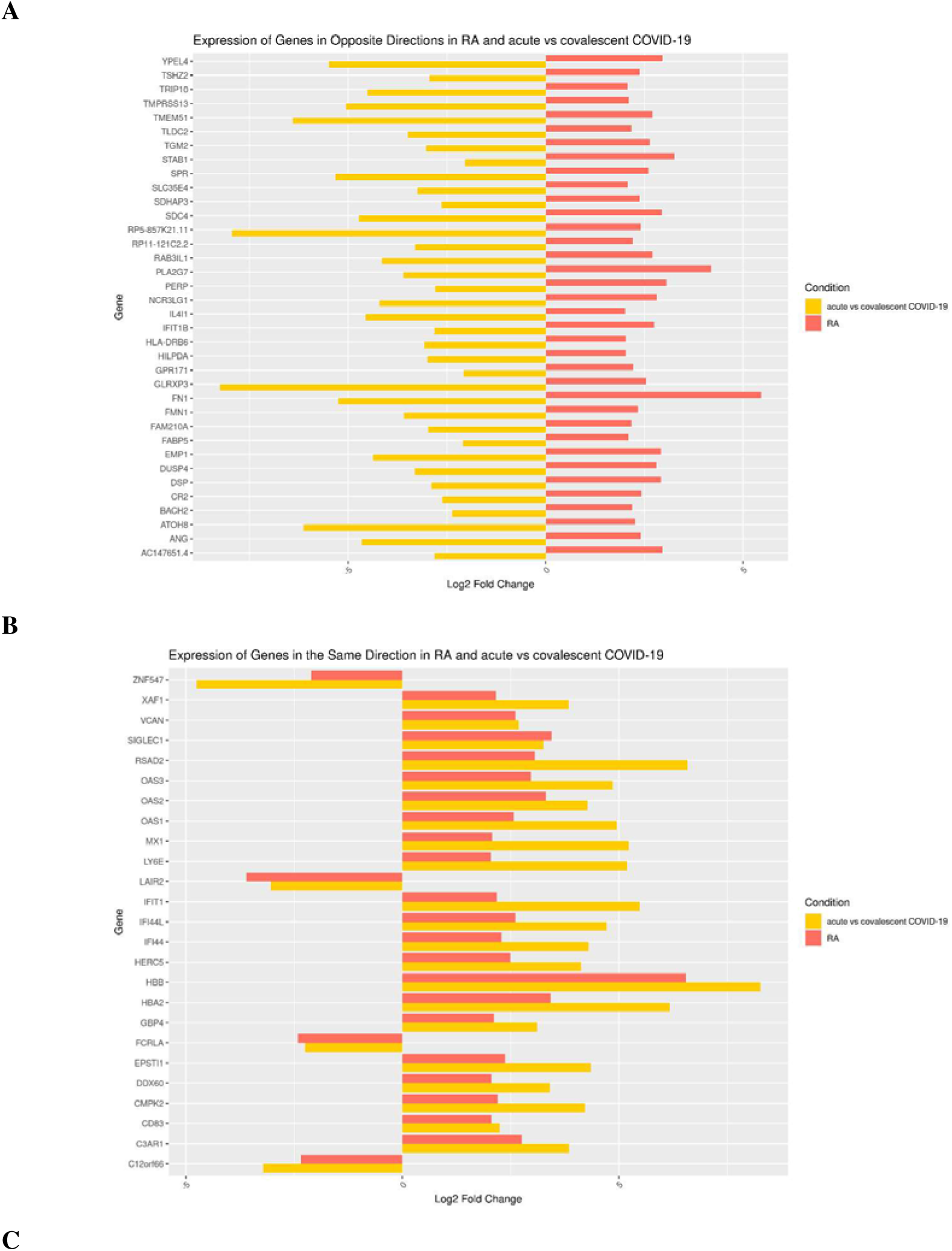
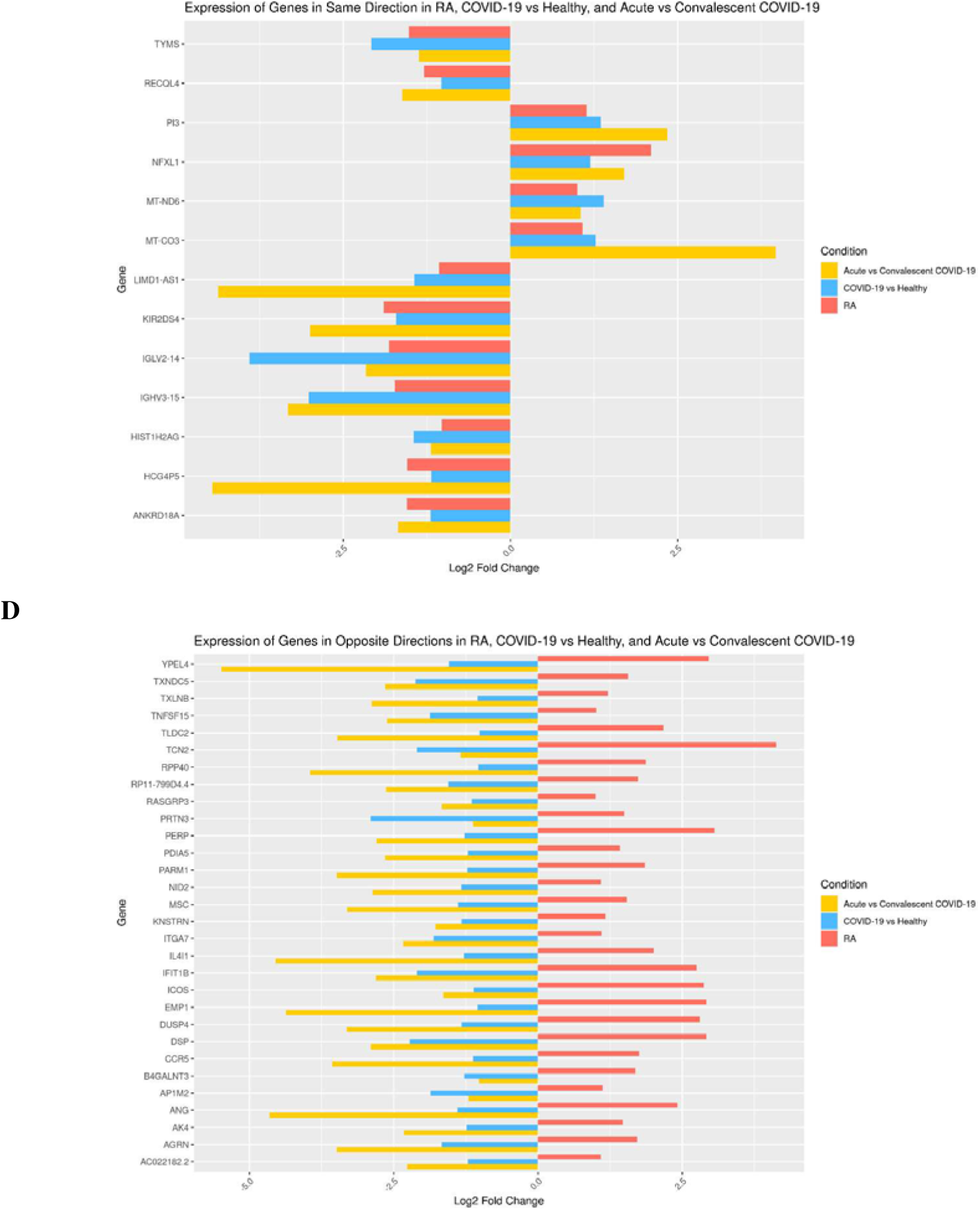
RA DEGs compared with COVID-19 cohorts 1 and 2. (A) Genes with opposite direction of change in RA and in cohort 2 (acute vs convalescent COVID-19). (B) Genes with the same direction of change in RA and cohort 2. (C) Genes with the same direction in RA, COVID-19 vs healthy (cohort 1) and acute vs convalescent COVID-19 (cohort 2). (D) Genes with opposite direction in RA relative to both COVID-19 comparisons. Bars show log2 fold change.

Notably, OAS1, OAS3, SIGLEC1, OAS2, MX1, RSAD2, IFIT1, IFI44, IFI44L, XAF1 and C3AR1 were among the genes changing in the same direction in RA and cohort 2 (Fig. 11B), whereas OAS1, OAS3, SIGLEC1, RSAD2, IFIT1, IFI44L, XAF1 and C3AR1 were downregulated in cohort 1 (Fig. 8).

## 4. Discussion

In RA neutrophils, differential expression and pathway analysis point to a prominent interferon-related and host-defence signature: interferon alpha and gamma responses were the top-ranked Hallmark gene sets, and GO enrichment of the 90 strongest DEGs featured type I interferon, antiviral and double-stranded RNA terms. This is notable because RA is not generally considered pathogen-driven, and it is consistent with the reported dual role of type I interferons in viral infection and autoimmunity [3], [11].

COVID-19 neutrophils in cohort 1 showed a different profile: most DEGs (383 of 420) were downregulated, and none of the Hallmark gene sets reached adjusted p<0.05. GO terms among the DEGs nevertheless overlapped with those found in RA; GO over-representation of a DEG set does not by itself indicate the direction of change. The overlap between the diseases at gene level was partial: 46 of the 90 RA DEGs moved in the same direction in COVID-19 without reaching significance, and 27 moved significantly in the opposite direction. These 27 include CAMK1G, SIGLEC1, OAS1 and OAS3, which are involved in immune-cell activation, immune signalling and antiviral immunity, respectively. Consistently, genes in the negative regulation of type I interferon-mediated signalling pathway were upregulated in RA and downregulated in COVID-19 cohort 1.

The second COVID-19 comparison qualifies this picture. Genes such as OAS1, OAS3 and SIGLEC1 changed in the same direction as RA in the acute-versus-convalescent comparison, whereas a different set of 30 genes was upregulated in RA and downregulated in both COVID-19 comparisons. The gene-level divergence between RA and COVID-19 therefore depends on the COVID-19 comparison used, and a gene should be considered distinct to RA only where it differs in both; on this criterion the 30 genes in Fig. 11D, rather than OAS1, OAS3 or SIGLEC1, are the more consistent candidates (CAMK1G was not among the genes displayed for cohort 2).

Published accounts describe severe COVID-19 as an acute, systemic process with neutrophilia, an increased neutrophil-to-lymphocyte ratio and cytokine-storm-associated tissue injury [7], whereas RA inflammation is chronic and mainly synovial [5], [10]. These are literature statements that our transcriptomic comparison did not test. The COVID-19 neutrophils in cohort 1 come from hospitalised patients described in the source studies [16], [17], so our findings should not be extrapolated to severe disease.

Epidemiological studies associate COVID-19 with an increased risk of RA and other autoimmune diseases [8], [9]. The partial overlap of neutrophil pathways found here is compatible with a role for shared immune dysregulation, but this analysis cannot establish that COVID-19 causes or worsens RA; this remains a hypothesis.

### Limitations

- Sample sizes are small. The RA count matrix includes technical replicates (the same library sequenced on two lanes), and RA samples from synovial fluid and peripheral blood were analysed together against healthy controls.
- Cohort 2 was assembled by merging two independent studies (severe versus healthy and convalescent versus healthy), so study-level differences may contribute to the results.
- DEGs were selected using p<0.05 with a log2 fold change threshold, and several enrichment results were displayed at an adjusted p<0.25.
- The findings are computational and have not been validated experimentally.

### Future directions

Priorities are (i) deeper study of shared immune pathways, especially type I interferon responses, and extension to other autoimmune diseases; (ii) experimental validation of candidate genes such as CAMK1G, SIGLEC1, OAS1 and OAS3 as biomarkers or therapeutic targets; (iii) longitudinal studies of people recovering from COVID-19 to assess the risk of RA and other autoimmune disorders; (iv) development of targeted therapies that modulate neutrophils; and (v) integration of proteomics, metabolomics and single-cell RNA sequencing.

## 5. Conclusion

Neutrophils from RA patients show a transcriptional programme enriched for interferon and antiviral-response pathways. Compared with COVID-19 neutrophils, RA shares part of this programme and diverges for a subset of genes, with the divergence depending on the COVID-19 comparison used. These results provide candidate genes and pathways for experimental validation and for understanding the relationship between COVID-19 and RA.

## Declarations

### Data availability

All datasets analysed are publicly available in GEO under the accessions listed in Table 1.

## Acknowledgements

The authors thank the faculty, lab members and staff of the Department of Computer Science, Jamia Millia Islamia, for support and access to computational resources.

## Funding, competing interests, author contributions, code availability

[to be completed by the authors].

## Supplementary Figures

**Supplementary Figure S1.**
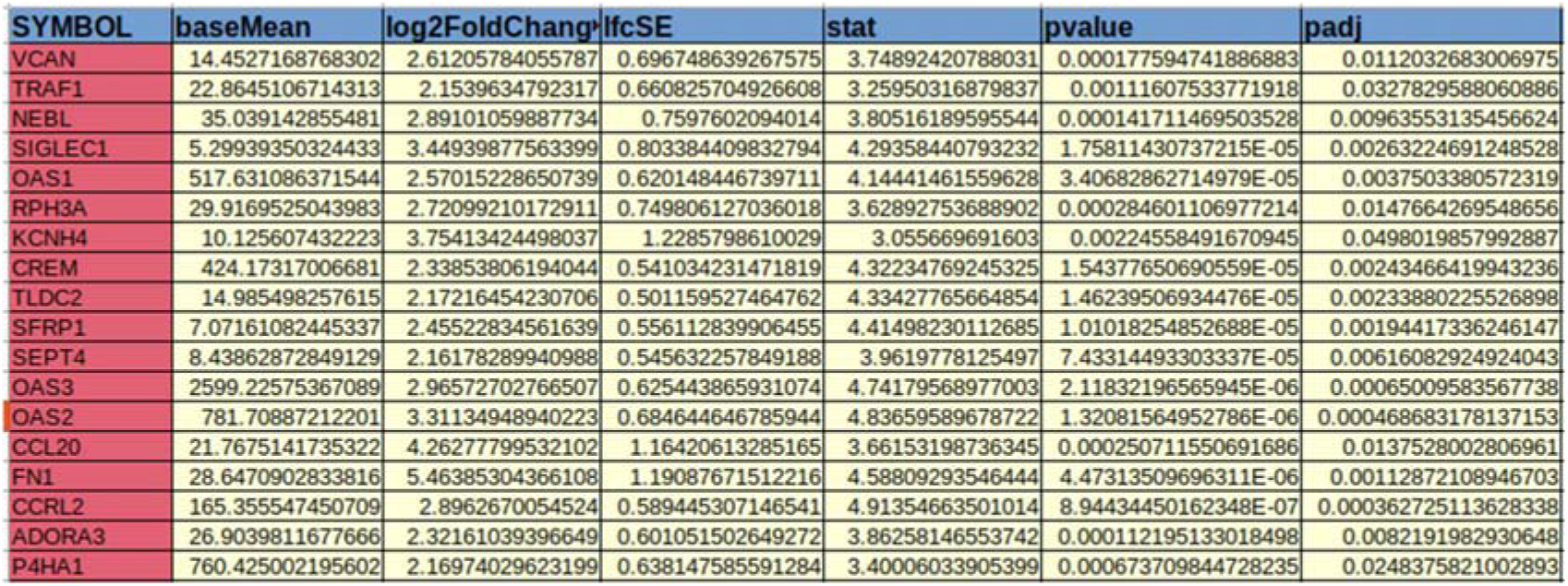
DEG output snippet for RA versus healthy control neutrophils (GSE116899). Columns show base mean, log2 fold change, standard error, test statistic, p-value and adjusted p-value.

**Supplementary Figure S2.**
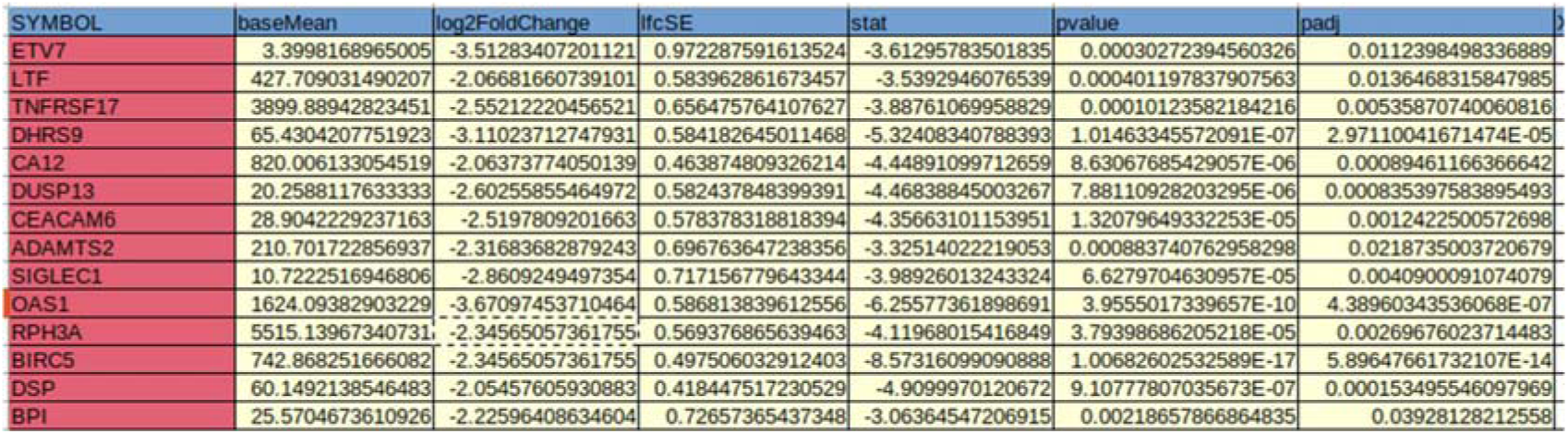
DEG output snippet for COVID-19 versus healthy control neutrophils (GSE212041). Columns as in Supplementary Fig. S1.

