## Supplementary figures and images for "Comparative Transcriptomic Analysis of Gene Expression Patterns in Neutrophils from Rheumatoid Arthritis and COVID-19"

### Supplemental Figure 1

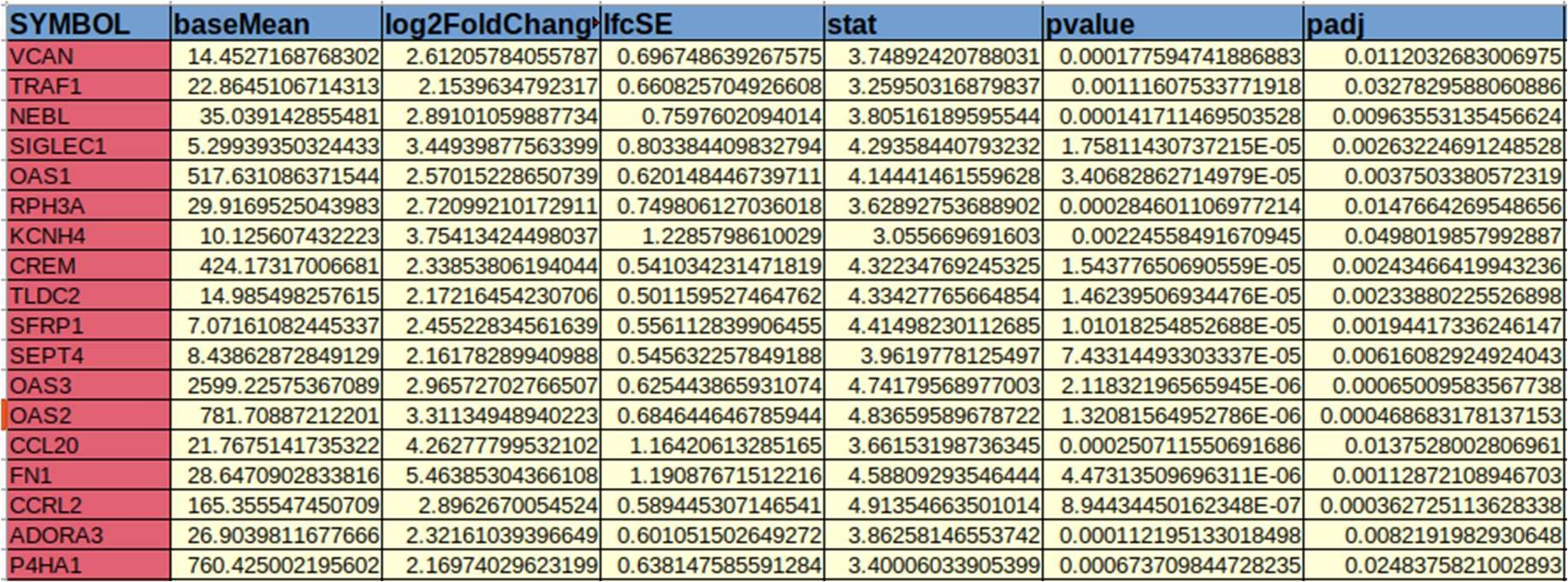

### Supplemental Figure 2

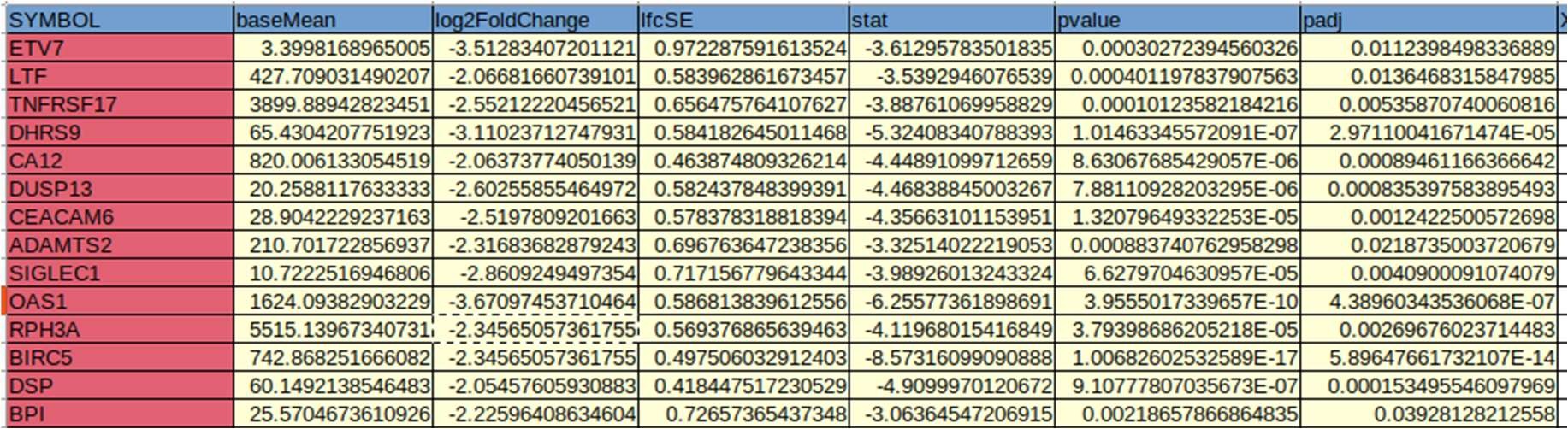
